# Long term conservation of DNA at ambient temperature. Implications for DNA data storage

**DOI:** 10.1101/2021.07.28.454193

**Authors:** Delphine Coudy, Marthe Colotte, Aurélie Luis, Sophie Tuffet, Jacques Bonnet

## Abstract

DNA conservation is central to many applications. This leads to an ever-increasing number of samples which are more and more difficult and costly to store or transport. A way to alleviate this problem is to develop procedures for storing samples at room temperature while maintaining their stability. A variety of commercial systems have been proposed but they fail to completely protect DNA from deleterious factors, mainly water. On the other side, Imagene company has developed a procedure for long-term conservation of biospecimen at room temperature based on the confinement of the samples under an anhydrous and anoxic atmosphere maintained inside hermetic capsules. The procedure has been validated by us and others for purified RNA, and DNA in buffy coat or white blood cells lysates, but a precise determination of purified DNA stability is still lacking. We used the Arrhenius law to determine the DNA degradation rate at room temperature. We found that extrapolation to 25 °C gave a degradation rate constant equivalent to about 1 cut/century/100 000 nucleotides, a stability several orders of magnitude larger than the current commercialized processes. Such a stability is fundamental for many applications such as the preservation of very large DNA molecules (particularly interesting in the context of genome sequencing) or oligonucleotides for DNA data storage. Capsules are also well suited for this latter application because of their high capacity. One can calculate that the 64 zettabytes of data produced in 2020 could be stored, standalone, for centuries, in about 20 kg of capsules.

## Introduction

Conservation of DNA, purified, in biospecimens or synthetic is a prerequisite to many applications, from biobanking, biodiversity preservation or molecular diagnostics to digital data storage (for reviews, see for instance [1,2,3]). This generates an ever-increasing number of samples which are more and more difficult and costly to store or transport. For reviews, see[4,5,6].

A way to alleviate, at least partially, this problem, is to develop procedures for storing samples at room temperature while maintaining their stability. Indeed, room temperature allows a standalone storage without energy costs. This implies to keep DNA away from environmental degradation factors, water, oxygen, ozone, and other atmospheric pollutants[7,8,9,10], water being by far the most deleterious element.

Many systems, often based on dehydration, have been used for room temperature storage of purified DNA: freeze-drying [11], inclusion in soluble matrices including liposomes, polymers such as silk [12] or pullulan [13] or adsorption on solid supports such as natural or treated cellulose [14,15,16]. Other procedures use encapsulation in sol-gel-based silica [17,18] or in silica nanoparticles [19,20], inclusion in salts [21] or layered double hybrids [22], dissolution in deep eutectic solvents [23] or ionic liquids [24]. As none of these procedures can totally protect DNA from atmosphere or moisture, other ways have been proposed: protection under a gold film [25] or encapsulation under an inert atmosphere in hermetic stainless-steel capsules, the DNAshells™ (Imagene SA, France) [6,26,27].

To demonstrate the real efficacy of a given preservation procedure one must estimate the DNA rate of degradation at room temperature (here, 25 °C) which is difficult because of the low degradation rate of dehydrated DNA in this condition. So, generally, one must rely on accelerated aging kinetics and extrapolation to room temperature by using Arrhenius equation. Such an approach has recently been used by Grass et al [28] and Organick et al [29], to compare some of these procedures in the context of DNA data storage. Among the tested procedures, DNA encapsulated in DNAshell™ did not give reliable rates of degradation because these were too low.

Here we report an Arrhenius analysis for purified DNA stored in DNAshell complementing these previous studies and exemplifying the high stability of DNA when stored under inert atmosphere.

## Material and methods

### DNA preparation

DNA was extracted from blood collected on EDTA, following the Puregene protocol (Gentra, Qiagen, Hilden Germany) and resuspended in 10 mM Tris-HCl, 1 mM EDTA, pH 8 and stored at 4 °C.

### Ethics statement

The data regarding DNA stability presented in this study relate to projects that have been formally approved by the “*Comité de protection des personnes Sud Ouest et Outre Mer III”º*, including use of blood and blood-derived samples. “*L’Etablissement français du sang*” (EFS, France) is a French national establishment that is authorized to collect blood samples from adult volunteer donors for both therapeutic and non-therapeutic uses. The donations were collected in accordance with the French blood donation regulations and ethics and with the French Public Health Code (art L1221-1). Blood samples were anonymized according to the French Blood Center (EFS) procedure. Volunteer donors signed written informed consents before blood collection. EFS authorized Imagene to perform this study and provided de-identified blood and blood-derived samples for non-therapeutic use.

### DNA encapsulation

DNA encapsulation was realized as previously described. Briefly, the DNA solutions (700 ng in 10 µL) were aliquoted in glass insert fitted in open stainless-steel capsules (DNAshells). The samples were dried under vacuum and left overnight in a glove box under an anoxic and anhydrous argon/helium atmosphere for further desiccation. Then, caps were added and sealed by laser welding. Finally, the DNAshells were checked for leakage by mass spectrometry [27,30].

### DNA accelerated degradation studies

DNAshells were heated in a Thermoblock at 100 °C, 110 ° C,120 °C, 130 °C and 140°C. At each time point of the kinetics, 2 or 3 capsules were retrieved, and stored at -20°C. For analysis, the capsules were opened, the DNA s were rehydrated with 20 µL of water and immediately analyzed by electrophoresis or stored at -20 °C for qPCR.

### Measure of DNA degradation rates by qPCR with two different sizes amplicons

Two amplicons of 1064 bp and a 93 bp of the TAF1L gene (TATA-box binding protein associated factor 1 like, Gene ID: 138474) were targeted. For both systems, PCR cycles were as follow: 10 min at 95 °C then 45 cycles of (15 s at 95 °C; 15 s at 60 °C; 60 s at 72 °C). The primers sequences were:

For-5’ agactcggacagcgaggaa/ Rev-5’ cggagacacccagcatatca for the 1064 pb fragment and

For-5’ tgcaggcacttgagaacaac / Rev-5’ aaccctgtcttgtccgaatg for the 93 pb.

They were produced by Eurogenetec, Les Ulis, France. After rehydration, for each sample and each amplicon, we used three ten-fold dilutions to estimate the PCR efficiency. These samples and their dilutions were analyzed independently and defined as “standards”.

To determine the degradation rates, we used a previously developed model [9] based on qPCR amplification of two amplicons (1 and 2) of different sizes (L_1_ and L_2_) to measure the number of cuts per nucleotide (or the probability of breakage at a given position). Assuming a random breakage mechanism, the probability of breakage at a given position is:

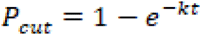

and the probability of this position remaining unbroken is:

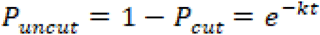

From the model:

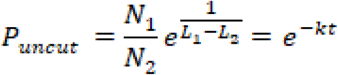

So, for each temperature, T, a graph of

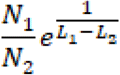

versus time gave us k_T_ by curve fitting.

This method is more reliable than one-sized qPCR because it does not rely on initial amounts of sample and, so, avoids errors due to losses upon recovery.

## Results and discussion

### Measure of DNA degradation rates by qPCR with two different sizes amplicons

The qPCR analyses gave the number of amplifiable copies N_1064_ and N_93_ for each time point (t) and temperature (T) and we plotted P_uncut_ versus time to obtain k_T for_ each temperature. (Fig 1)

**Fig 1.**
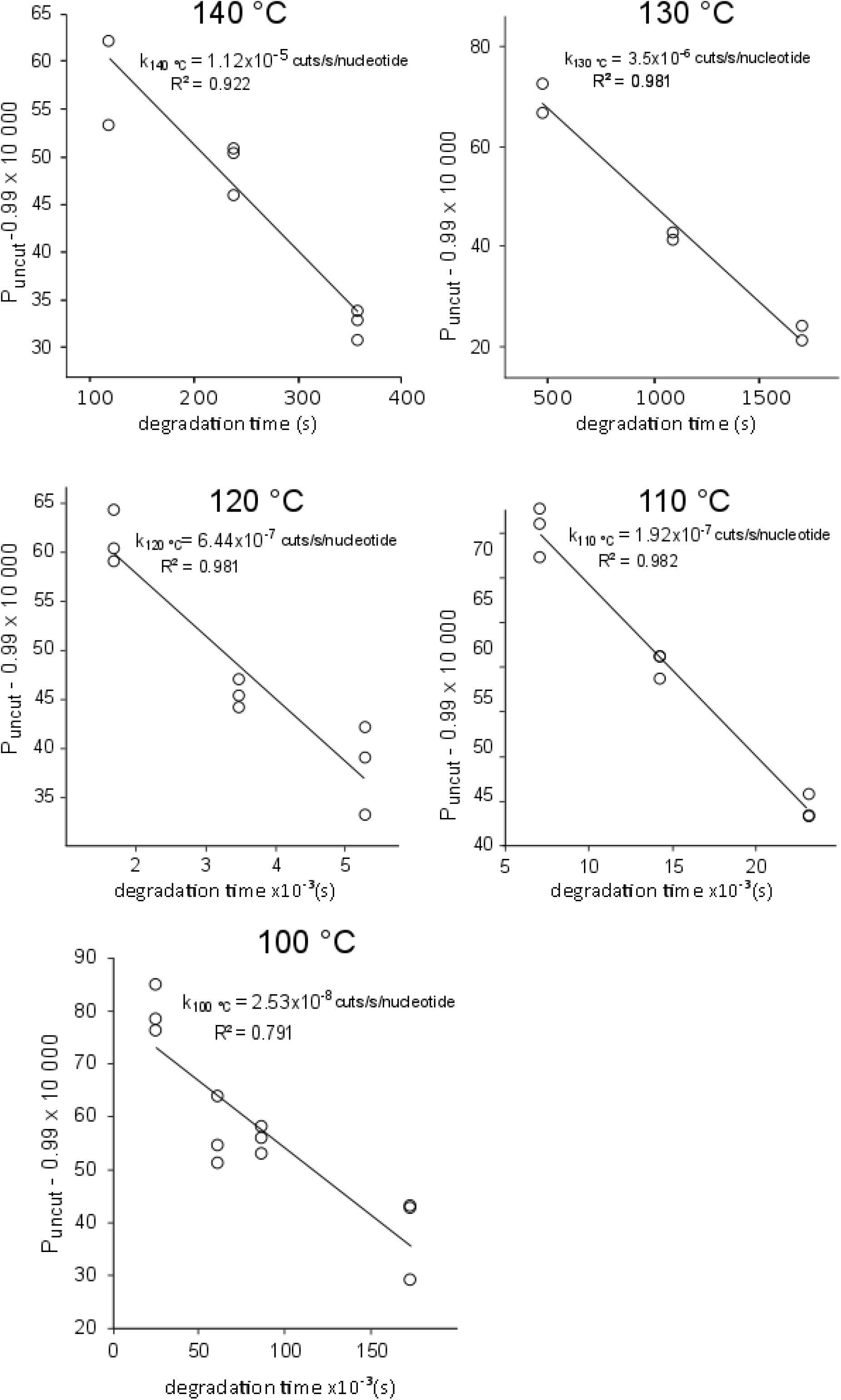
Degradation kinetics of DNA stored in DNAshells. For each time point, the proportion of intact nucleotide position, P_uncut_, was calculated from the numbers N_1064_ and N_93_ of amplifiable copies of the 1064 pb and 93 pb amplicons. These values were plotted as a function of the degradation time at the indicated temperatures. The lines are the fit to the data points by Microsoft Excel software.

Then, we plotted the logarithm of k_T_ as a function of the reverse of the absolute temperature (T) (Fig 2).

**Fig 2.**
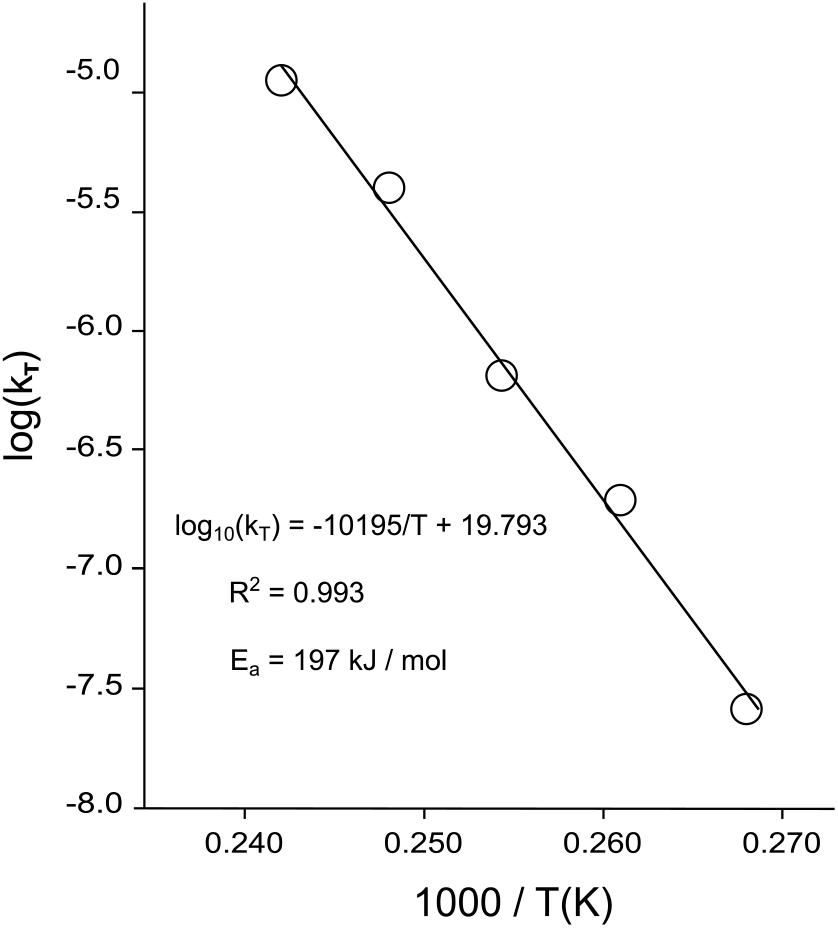
Arrhenius plot for DNA degradation in DNAshells. The degradation rate constants, k, were plotted as a function of the reverse of the absolute temperature T.

This plot showed that the degradation rate followed the Arrhenius law with an activation energy of 197 kJ/mol. This is comparable to the 163 kJ/mol to 188kJ/mol previously found for desiccated plasmid DNA [7] and to about the 155 kJ/mol for DNA stored in silica nanoparticles, FTA paper or DNAstable matrix [28]. This is significantly higher than the 100 kJ/mol to 121 kJ/mol found for degradation of double strand DNA in solution (reviewed in [7]).

The Arrhenius law also made it possible to extrapolate the degradation rate at 25 °C. This gave a degradation rate constant of 3.82×10^−15^ cuts /s/nucleotide, equivalent to about 1 cut/century/100 000 nucleotides or 38 000 years of half-life for a 150-nucleotide long DNA fragment (we chose this size for an immediate comparison with the previous works [28,29]).

This allows to calculate the time necessary for a DNA molecule to degrade down to 25 nucleotides, the average length which is the current limit for the sequencing of degraded DNA [31] using the formula:

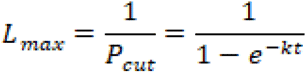

This gives 1,070,000 years, provided the preservation conditions are maintained.

Fig 3 compares the half-lives of a 150 nucleotides long DNA fragment stored in various conditions at 25 °C.

**Fig 3.**
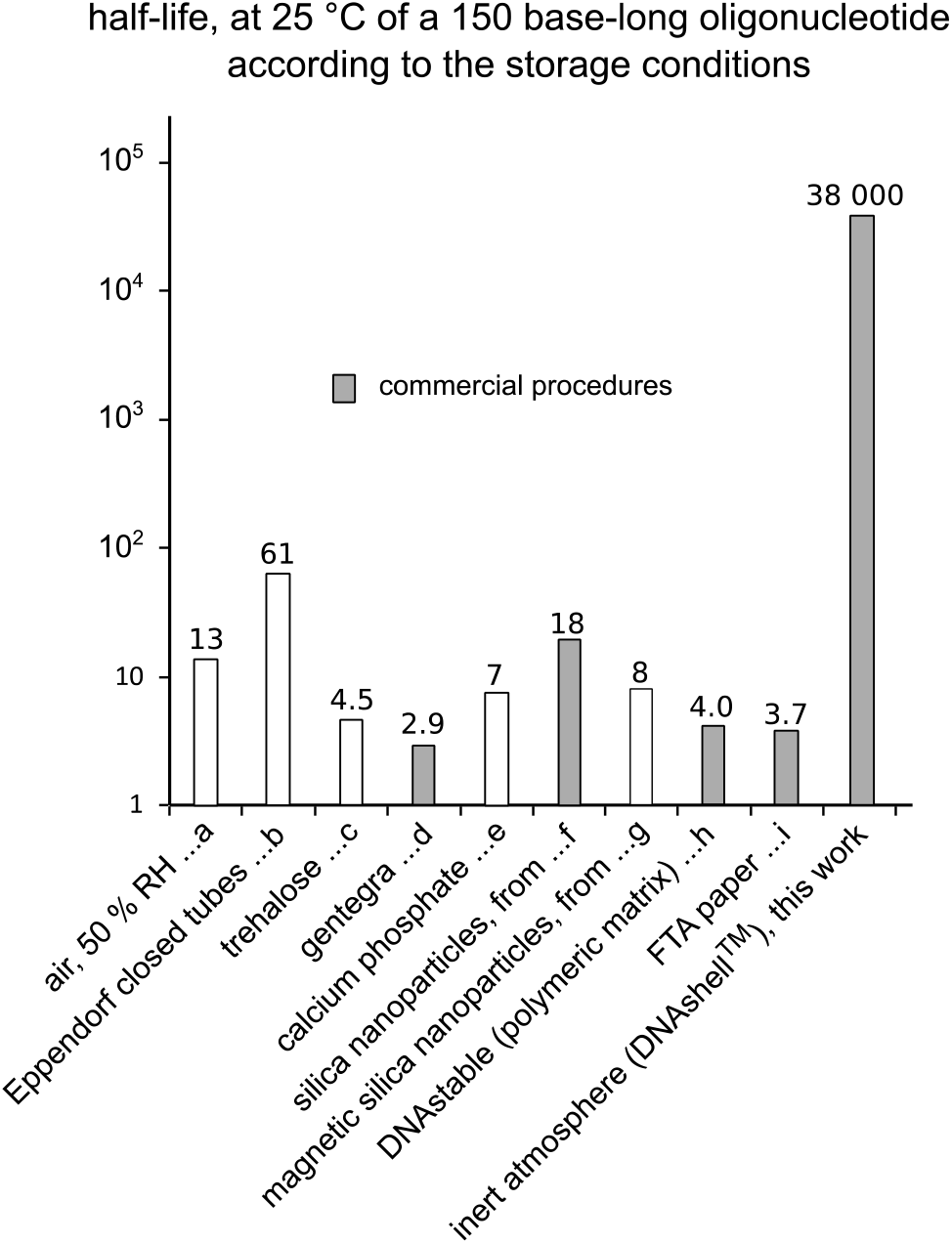
Half-lives of a 150 nucleotides long DNA fragment stored in various conditions. The half-life of a DNA sample left unprotected from the atmosphere at room temperature (a) or in an Eppendorf closed tube (b) has been calculated from our previous work [7]. The one of a sample encapsulated in silica nanoparticles (g), deposited on FTA card paper (i) or included in Biomatrica DNAstable (h) has been estimated from [28] (Fig 2B). The half-life of DNA dried with calcium phosphate (e) and encapsulated in magnetic silica nanoparticles (f) have been estimated, respectively from [29] (Fig 3b) and [19] (Fig 5) assuming an exponential decay and an activation energy of 155 kJ/molThe half-life for DNA stored in Gentegra (d) or trehalose (c) was taken from [29] (Fig 2b). In grey: current commercialized procedures.

So, it appears that the DNA stability at room temperature (25 ° C) is over three orders of magnitude higher in DNAshells than in any of the other currently commercialized storage devices.

This is to be expected because, first, FTA paper, trehalose or calcium phosphate leaves the DNA samples directly exposed to the atmosphere. Second, likewise, the matrices coating DNA: Gentegra, DNA stable and trehalose being water soluble cannot either protect the sample from moisture. Finally, silica nanoparticles, while affording protection from atmosphere, still contain a certain amount of water [28].

It must be noticed that the experiments conducted here mainly detect chain breaks, so, other degradations events not preventing elongation by the polymerase could go undetected. However, this should not be a concern because these are dependent on water and much slower than depurination and chain breaks [32]. Moreover, Organick et al sequenced the DNA samples stored at 85 °C for 4 weeks without noticing any increase in error rates while there is a significant number of chain breaks [29].

As a conclusion, this procedure, allowing a standalone storage is well suited for long term preservation of DNA samples for whatever application, in particular the recently developing DNA data storage procedures. Our figures make it possible to give an estimation of the lifetime of the data stored that way. Indeed, according to a recent estimation by Organick et al [33], 10 is the lowest copy number necessary for a faithful data storage and retrieval. This means that if one starts with 20 copies, the data could faithfully be retrieved after 30 centuries of storage.

Another advantage is the volume of the capsule which can accommodate large amounts of DNA. With a 200 µL useful volume and a DNA density of 1.4 [34], a single capsule could store 0.28 g of DNA. According to [35] estimating at about 17 exabytes/g the data density in DNA, this corresponds to 4.76 exabytes of data per DNAshell, equivalent to 1.6×10^12^ files (assuming an average file size of 3 Mo). Of course, it may look difficult to recover a specific file among this mass of data, however, this seems possible as described recently by Tomek et al claiming that, by using a combination of N primers, it could be possible to select a given file in a population of 27 999 ^N^ files [36].

So, according to these figures, the 64 zettabytes of data produced in 2020 [37] could theoretically be coded in 3,765 g of DNA which could be stored in 13,445 capsules packed in a suitcase weighing 21 kg.

This procedure could also allow the long-term room temperature preservation of very large DNA molecules which is particularly interesting in the context of genome sequencing, as a recent paper by Nurk et al described for the first time the sequencing of a complete human genome thanks to the use of very long DNA stretches [38].

## Acknowledgement

We thank Régine Gandoin for editing the manuscript.

## Competing interests

The authors have conflicts of interest in relation to the submitted work: D. Coudy, A. Luis, S. Tuffet and M. Colotte are employees of Imagene Company; S. Tuffet is CEO and share-holder of Imagene Company; J. Bonnet is a shareholder and a consultant of Imagene company.

## Author Contributions

**Conceptualization:** Delphine Coudy, Marthe Colotte, Aurélie Luis, Sophie Tuffet, Jacques Bonnet.

**Formal analysis:** Delphine Coudy, Marthe Colotte, Jacques Bonnet.

**Funding acquisition:** Sophie Tuffet

**Investigation** Delphine Coudy, Marthe Colotte.

**Methodology:** Delphine Coudy, Marthe Colotte, Aurélie Luis, Jacques Bonnet.

**Project administration:** Delphine Coudy, Marthe Colotte.

**Supervision:** Sophie Tuffet, Jacques Bonnet.

**Validation:** Delphine Coudy, Marthe Colotte, Aurélie Luis, Sophie Tuffet, Jacques Bonnet.

**Visualization:** Delphine Coudy, Marthe Colotte, Jacques Bonnet

**Writing – original draft:** Jacques Bonnet.

**Writing – review & editing** Delphine Coudy, Marthe Colotte, Aurélie Luis, Sophie Tuffet, Jacques Bonnet.

## References

1. Ceze L, Nivala J, Strauss K. Molecular digital data storage using DNA. Nature Reviews Genetics. 2019; 20(8): 456–466. doi: 10.1038/s41576-019-0125-3

2. Lim CK, Nirantar S, Yew WS, Poh CL. Novel Modalities in DNA Data Storage. Trends Biotechnol. 2021. doi: 10.1016/j.tibtech.2020.12.008

3. Dong Y, Sun F, Ping Z, Ouyang Q, Qian L. DNA storage: research landscape and future prospects. National Science Review. 2020; 7(6): 1092–1107. doi: 10.1093/nsr/nwaa007

4. Lou JJ, Mirsadraei L, Sanchez DE, Wilson RW, Shabihkhani M, Lucey GM, et al. A review of room temperature storage of biospecimen tissue and nucleic acids for anatomic pathology laboratories and biorepositories. Clinical Biochemistry. 2014; 47(4-5): 267–273. doi: 10.1016/j.clinbiochem.2013.12.011

5. Muller R, Betsou F, Barnes MG, Harding K, Bonnet J, Kofanova O, et al. Preservation of Biospecimens at Ambient Temperature: Special Focus on Nucleic Acids and Opportunities for the Biobanking Community. Biopreserv Biobank. 2016; 14(2): 89–98. doi: 10.1089/bio.2015.0022

6. Washetine K, Kara-Borni M, Heeke S, Bonnetaud C, Felix JM, Ribeyre L, et al. Ensuring the Safety and Security of Frozen Lung Cancer Tissue Collections through the Encapsulation of Dried DNA. Cancers (Basel). 2018; 10(6). doi: doi:10.3390/cancers10060195

7. Bonnet J, Colotte M, Coudy D, Couallier V, Portier J, Morin B, et al. Chain and conformation stability of solid-state DNA: implications for room temperature storage. Nucleic Acids Res. 2010; 38(5): 1531–1546. doi: 10.1093/nar/gkp1060

8. Cataldo F. DNA degradation with ozone. Int J Biol Macromol. 2006; 38(3-5): 248–254. doi: 10.1016/j.ijbiomac.2006.02.029

9. Colotte M, Couallier V, Tuffet S, Bonnet J. Simultaneous assessment of average fragment size and amount in minute samples of degraded DNA. Anal Biochem. 2009; 388345– 347. doi: 10.1016/j.ab.2009.02.003

10. Pacini S, Giovannelli L, Gulisano M, Peruzzi B, Polli G, Boddi V, et al. Association between atmospheric ozone levels and damage to human nasal mucosa in Florence, Italy. Environ Mol Mutagen. 2003; 42(3): 127–135. doi: 10.1002/em.10188 [doi]

11. Vilenchik MM. Studies of DNA damage and repair of thermal-and radiation-induced lesions in human cells. Int J Radiat Biol. 1989; 56(5): 685-689. doi:

12. Liu Y, Zheng Z, Gong H, Liu M, Guo S, Li G, et al. DNA preservation in silk. Biomater Sci. 2017. doi: 10.1039/c6bm00741d [doi]

13. Jahanshahi-Anbuhi S, Pennings K, Leung V, Liu M, Carrasquilla C, Kannan B, et al. Pullulan encapsulation of labile biomolecules to give stable bioassay tablets. Angew Chem Int Ed Engl. 2014; 53(24): 6155–6158. doi: 10.1002/anie.201403222 [doi]

14. Kline MC, Butts ELR, Hill CR, Coble MD, Duewer DL, Butler JM. The new Standard Reference Material® 2391c: PCR-based DNA profiling standard. Forensic Science International: Genetics Supplement Series. 2011; 3(1): e355-e356. doi:

15. Owens CB, Szalanski AL. Filter paper for preservation, storage, and distribution of insect and pathogen DNA samples. J Med Entomol. 2005; 42(4): 709-711. doi:

16. Smith LM, Burgoyne LA. Collecting, archiving and processing DNA from wildlife samples using FTA® databasing paper. BMC Ecology. 2004; 4(1): 4. doi: 10.1186/1472-6785-4-4

17. Pierre A, Bonnet J, Vekris A, Portier J. Encapsulation of deoxyribonucleic acid molecules in silica and hybrid organic-silica gels. J Mater Sci Mater Med. 2001; 12(1): 51-55. doi: 319855 [pii]

18. Kapusuz D, Durucan C. Exploring encapsulation mechanism of DNA and mononucleotides in sol-gel derived silica. J Biomater Appl. 2017; 885328217713104. doi: 10.1177/0885328217713104 [doi]

19. Chen WD, Kohll AX, Nguyen BH, Koch J, Heckel R, Stark WJ, et al. Combining Data Longevity with High Storage Capacity Layer-by-Layer DNA Encapsulated in Magnetic Nanoparticles. Advanced Functional Materials. 2019; 0(0): 1901672. doi: 10.1002/adfm.201901672

20. Chen WD, Kohll AX, Nguyen BH, Koch J, Heckel R, Stark WJ, et al. Combining Data Longevity with High Storage Capacity—Layer-by-Layer DNA Encapsulated in Magnetic Nanoparticles. Advanced Functional Materials. 2019; 29(28): 1901672. doi: 10.1002/adfm.201901672

21. Kohll AX, Antkowiak PL, Chen WD, Nguyen BH, Stark WJ, Ceze L, et al. Stabilizing synthetic DNA for long-term data storage with earth alkaline salts. Chem Commun (Camb). 2020. doi: 10.1039/d0cc00222d [doi]

22. Choy J-H, Kwak S-Y, Park J-S, Jeong Y-J, Portier J. Intercalative Nanohybrids of Nucleoside Monophosphates and DNA in Layered Metal Hydroxide. Journal of the American Chemical Society. 1999; 121(6): 1399–1400. doi: 10.1021/ja981823f

23. Rizana Y, Haslina Ahmad, Mohd Basyarud0din, Abdul Rahman MB. Studies of interaction between tetrabutylammonium bromide based deep eutectic solvent and dna using fluorescence quenching method and circular dichroism spectroscopy. Malaysian Journal of Analytical Sciences. 2016; 20(6): 1233–1240. doi: 10.17576/mjas-2016-2006-01

24. Singh N, Sharma M, Mondal D, Pereira MM, Prasad K. Very high concentration solubility and long term stability of DNA in an ammonium-based ionic liquid: a suitable media for nucleic acid packaging and preservation. ACS Sustainable Chemistry & Engineering. 2017. doi: 10.1021/acssuschemeng.6b02842

25. Jiang J (2008) Gold film protected biomolecules in micro-fabrication, Biopatterning and DNA storage. Thesis, the National University of Singapore. Available from http://scholarbank.nus.edu.sg/handle/10635/19232.

26. Clermont D, Santoni S, Saker S, Gomard M, Gardais E, Bizet C. Assessment of DNA encapsulation, a new room-temperature DNA storage method. Biopreserv Biobank. 2014; 12(3): 176–183. doi: 10.1089/bio.2013.0082

27. Colotte M, Coudy D, Tuffet S, Bonnet J. Adverse Effect of Air Exposure on the Stability of DNA Stored at Room Temperature. Biopreserv Biobank. 2011; 9(1): 47–50. doi: 10.1089/bio.2010.0028

28. Grass RN, Heckel R, Puddu M, Paunescu D, Stark WJ. Robust Chemical Preservation of Digital Information on DNA in Silica with Error-Correcting Codes. Angew Chem Int Ed Engl. 2015; 54(8): 2552–2555. doi: 10.1002/anie.201411378

29. Organick L, Nguyen BH, McAmis R, Chen WD, Kohll AX, Ang SD, et al. An Empirical Comparison of Preservation Methods for Synthetic DNA Data Storage. Small Methods. 2021; 5(5): 2001094. doi: 10.1002/smtd.202001094

30. Fabre AL, Luis A, Colotte M, Tuffet S, Bonnet J. High DNA stability in white blood cells and buffy coat lysates stored at ambient temperature under anoxic and anhydrous atmosphere. PLoS One. 2017; 12(11): e0188547. doi: 10.1371/journal.pone.0188547

31. Xavier C, Eduardoff M, Bertoglio B, Amory C, Berger C, Casas-Vargas A, et al. Evaluation of DNA Extraction Methods Developed for Forensic and Ancient DNA Applications Using Bone Samples of Different Age. Genes (Basel). 2021; 12(2). doi: 10.3390/genes12020146

32. Shen J-C, Rideout WM III, Jones PA. The rate of hydrolytic deamination of 5-methylcytosine in double-stranded DNA. Nucl Acids Res. 1994; 22(6): 972–976. doi: 10.1093/nar/22.6.972

33. Organick L, Chen Y-J, Ang SD, Lopez R, Strauss K, Ceze L. Experimental Assessment of PCR Specificity and Copy Number for Reliable Data Retrieval in DNA Storage. biorxiv [preprint] 2019; Available at https://www.biorxiv.org/content/10.1101/565150v1. doi: 10.1101/565150

34. Smialek MA, Jones NC, Hoffmann SrV, Mason NJ. Measuring the density of DNA films using ultraviolet-visible interferometry. Physical Review E. 2013; 87(6): 060701. doi: 10.1103/PhysRevE.87.060701

35. Organick L, Chen YJ, Dumas Ang S, Lopez R, Liu X, Strauss K, et al. Probing the physical limits of reliable DNA data retrieval. Nat Commun. 2020; 11(1): 616. doi: 10.1038/s41467-020-14319-8

36. Tomek KJ, Volkel K, Simpson A, Hass AG, Indermaur EW, Tuck JM, et al. Driving the Scalability of DNA-Based Information Storage Systems. ACS Synthetic Biology. 2019. doi: 10.1021/acssynbio.9b00100

37. Consortium INSIC 2019. Available at http://www.insic.org/wp-content/uploads/2019/07/INSIC-Technology-Roadmap-2019.pdf.

38. Nurk S, Koren S, Rhie A, Rautiainen M, Bzikadze AV, Mikheenko A, et al. The complete sequence of a human genome. bioRxiv. 2021; 2021.2005.2026.445798. doi: 10.1101/2021.05.26.445798

